# No effect of administration of unacylated ghrelin on subcutaneous PC3 xenograft growth in a *Rag1*^−/−^ mouse model of metabolic dysfunction

**DOI:** 10.1101/328351

**Authors:** Michelle L. Maugham, Inge Seim, Patrick B. Thomas, Gabrielle J. Crisp, Esha T. Shah, Adrian C. Herington, Kristy A. Brown, Laura S. Gregory, Colleen C. Nelson, Penny L. Jeffery, Lisa K. Chopin

## Abstract

Ghrelin is a peptide hormone which, when acylated, regulates appetite, energy balance and a range of other biological processes. Ghrelin predominately circulates in its unacylated form (unacylated ghrelin; UAG). UAG has a number of functions independent of acylated ghrelin, including modulation of metabolic parameters and cancer progression. UAG has also been postulated to antagonise some of the metabolic effects of acyl-ghrelin, including its effects on glucose and insulin regulation. In this study, *Rag1^−/−^* mice with high-fat diet-induced obesity and hyperinsulinaemia were subcutaneously implanted with PC3 prostate cancer xenografts to investigate the effect of UAG treatment on metabolic parameters and xenograft growth. Daily intraperitoneal injection of 100 μg/kg UAG had no effect on xenograft tumour growth in mice fed normal rodent chow or 23% high-fat diet. UAG significantly improved glucose tolerance in host *Rag1^−/−^* mice on a high-fat diet, but did not significantly improve other metabolic parameters. We hypothesise that UAG is not likely to be an effective treatment for prostate cancer, with or without associated metabolic syndrome.

**Conflict of interest:** The authors declare no conflict of interest.

## Introduction

The peptide hormone ghrelin is a circulating appetite-stimulating hormone which regulates a number of other biological processes [1–3], including metabolism and energy balance [1–4], and diseases such as cancer [5]. Ghrelin acts via its cognate receptor, the growth hormone secretagogue receptor 1a (GHSR1a), a G protein-coupled receptor [6], and one or more unknown alternative receptors [7–10]. In order to activate GHSR1a at physiological concentrations, ghrelin must be acylated at its third residue, a serine [11, 12], by the enzyme ghrelin *O*-acyl transferase (GOAT) [11, 12].

The major circulating form of ghrelin is its unmodified form, unacylated ghrelin (UAG). UAG, which does not directly stimulate feeding [5], was initially considered to be functionally inactive, but is now appreciated to bind to and activate a distinct, unknown receptor [4, 13–18] and have a number of functions [19–22]. UAG plays roles in the regulation of glucose and energy balance and has effects on cell proliferation [19–23]. Importantly, it may oppose some of the effects of acyl-ghrelin [16, 24–26] preventing the rise in circulating glucose and insulin associated with acyl-ghrelin administration in rodents [22, 27]. From these studies, it is apparent that UAG is an endocrine hormone in its own right [20]. UAG and the truncated, cyclised UAG analogue AZP-531 prevented the development of pre-diabetes in C57BL/6 mice fed a high-fat diet for two weeks, highlighting a potential of unacylated forms of ghrelin as treatments for metabolic syndrome [27]. In human trials, UAG had similar effects, improving glycaemic control and insulin sensitivity in patients with type 2 diabetes mellitus [28] and improving glucose handling and reducing free fatty acids in healthy subjects when administered overnight as a continuous infusion [29]. AZP-531 also had beneficial effects on glucose balance and led to weight loss in patients with type 2 diabetes mellitus in a phase I clinical trial [30]. Similar benefits have been observed in patients with Prader-Willi syndrome, a genetic disorder associated with hyperghrelinaemia and obesity [31].

Close to two decades of work has firmly established a role for the ghrelin axis in cancer [5, 32–35]. This includes prostate cancer, a classical endocrine-related cancer and the most commonly diagnosed cancer in American men after skin cancer [36], where acyl-ghrelin increases cell proliferation and migration [5, 14, 37–46]. UAG also has functional effects in several cancers, including prostate cancer [5, 32–34, 43]. In the PC3 prostate cancer cell line, UAG has a biphasic effect, reducing cell proliferation at supraphysiological levels (10nM-1μM) [14].

Studies investigating the role of UAG in prostate cancer have been limited to *in vitro* experiments. *In vivo* studies are required, however, as obesity and overweight and co-morbidities, including hyperinsulinaemia, are now recognised as critical risk factors for numerous cancers [47–49]. These include cancer types with high-prevalence and mortality, such as tumours of the prostate, endometrium, breast, and gastrointestinal system [47–55]. Obesity and increased body mass have been associated with increased risk of advanced prostate cancer, more aggressive and high-grade disease and increased risk of death from prostate cancer [56–59]. Castration-resistant prostate cancer (CRPC) occurs when prostate cancer recurs after remission from androgen-targeted therapies (ATT) [60]. Treatments for CRPC are limited and this stage of the disease often results in the formation of painful, metastatic bone lesions and associated morbidity and mortality [61–63]. Metabolic syndrome and hyperinsulinaemia are common side effects of ATT [64, 65] and may also further accelerate the progression to CRPC [48, 58, 66–68]. As UAG reduces prostate cancer proliferation *in vitro* [14] and has potential beneficial metabolic effects *in vivo*, we examined the effect of UAG in our model of metabolic dysfunction: *Rag1^−/−^* mice fed a high-fat diet, with subcutaneous prostate cancer cell line xenografts [69].

## Materials and Methods

### Cell Culture

Human prostate cancer cell lines were obtained from the American Type Culture Collection (ATCC, Manassas, VA, USA). The PC3 prostate cancer cell line was cultured in Roswell Park Memorial Institute 1640 medium (RPMI-1640) and supplemented with 10% (v/v) Fetal Calf serum (FCS) (Thermo Fisher Scientific, Waltham, MA, USA), 50 units/ml penicillin, and 100 μg/mL streptomycin (Thermo Fisher Scientific). Cells were tested negative for *Mycoplasma*.

### Hyperinsulinaemic *Rag1^−/−^* mouse model treated with unacylated ghrelin (UAG)

To determine the metabolic effect of UAG in an engraftable mouse model of hyperinsulinaemia [69], male recombination-activation gene deficient mice (B6.SVJ129-*Rag1^tm1Bal^*/Arc; *Rag1^−/−^*) (Jackson Laboratories, supplied by Animal Resource Centre, Murdoch, WA, Australia) were weaned onto a diet of low-fat normal chow (LFD) or a Western-style, high-fat diet (HFD; 23% fat, SF04-027, Specialty Feeds, WA) [69]. After two weeks on the diet, mice were subcutaneously injected into the left flank with 1×10^6^ PC3 cells diluted 1:1 in growth factor reduced, phenol red-free Matrigel (Corning, NY, USA). Tumours were allowed to grow until a volume of approximately 50-100 mm^3^ was reached, when mice were randomly divided into two experimental groups. Mice then received daily intraperitoneal injections of 100 μg/kg UAG (Mimotopes, Mulgrave, Vic, Australia) (**n*=6* HFD, *n*=10 LFD) (a dose previously determined to inhibit breast cancer growth *in vivo* [8]) or phosphate buffered saline (PBS) control (*n*=8 HFD, *n*=10 LFD) for 16 days. Tumour volume was calculated by measuring subcutaneous tumour length and width twice weekly using digital calipers (ProSciTech, Kirwan, QLD, Australia). Tumour volume was calculated using the equation ‘tumour volume = (width × length^2^)/2’ [70]. Bodyweight was measured twice weekly. At endpoint, tumours and adipose tissue (epididymal fat pad and interscapular brown adipose tissue) were excised and weighed. Fasting blood glucose was measured at endpoint and blood was collected by cardiac puncture for serum biochemical measurements. Surrogate indices of insulin resistance, insulin sensitivity, and steady state β-cell function were determined using the homeostatic model for assessment calculator (HOMA2), available from the Oxford Centre for Diabetes, Endocrinology and Metabolism [71], using measured fasting glucose and insulin levels.

### Statistics

Statistical analyses were performed using GraphPad Prism v.6.01 software (GraphPad Software, Inc., San Diego, CA). Kruskal-Wallis (three or more groups) and Mann-Whitney *U*-test (two groups) tests used for non-normally distributed data, while a two-way ANOVA with Tukey’s post-hoc test used for normally distributed data. *P* ≤ 0.05 was considered to be statistically significant.

## Results

### No effect of intraperitoneal administration of UAG on PC3 xenograft growth in obese, hyperinsulinaemic *Rag1^−/−^* mice

No significant differences in tumour volume over the treatment period or tumour weight (*P*=0.57) and volume at endpoint (*P*=0.55) were observed between UAG-treated and untreated obese mice (14 days of treatment) (Mann-Whitney test, Fig. 1A-C). An increase in insulin sensitivity (*P*=0.70) and reduction in body weight (*P*=0.08), epididymal fat pad weight (*P*=0.26), interscapular brown adipose tissue weight (*P*=0.12), fasting blood glucose (*P*=0.50), fasting blood insulin (*P*=0.90), insulin resistance (*P*=0.70), and steady-state P-cell function (*P*=0.22) was observed in the UAG treatment group in HFD-fed *Rag1^−/−^* mice-however, these changes were not statistically significant (Fig. 1D-L). There was a significant difference in blood glucose at 30 minutes following glucose challenge in HFD-fed UAG-treated mice compared to PBS controls at endpoint (after 16 days of treatment) (21.6 ± 1.2mM, *n*=6 vs 25.4 ± 1.9mM, *n*=5, *P*=0.02, two-way ANOVA with post-hoc test, Fig. 1G), however, this was not observed at other time points, suggesting no major change in glucose tolerance. There was no significant difference in fasting blood glucose (*P*=0.50), blood insulin concentration (*P* = 0.90, Mann-Whitney test, Fig. 1I), insulin resistance (*P* = 0.70, Mann-Whitney test, Fig. 1J), or insulin sensitivity (**P*=0.70*, Mann-Whitney test, Fig. 1L).

**Fig. 1.**
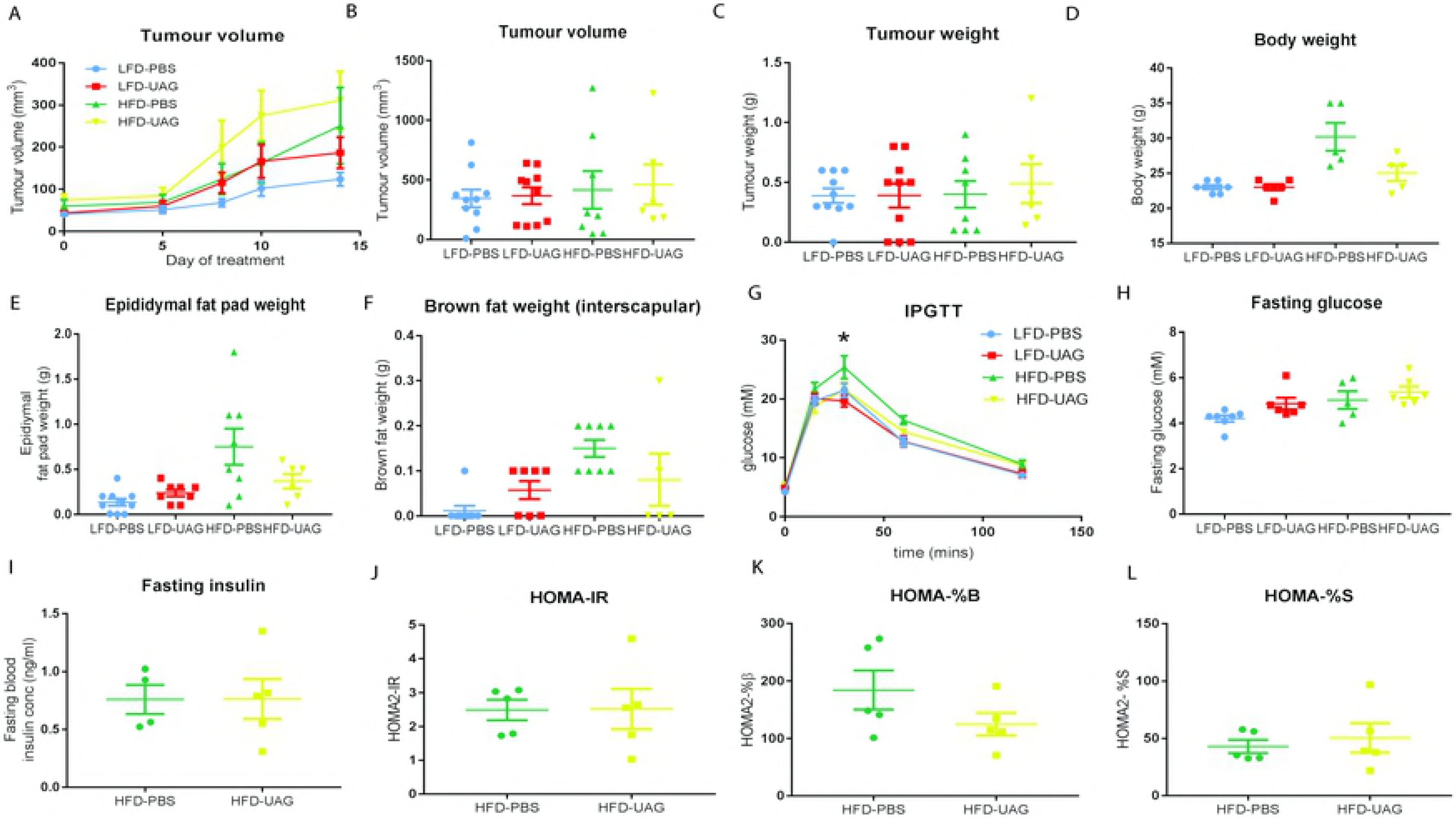
Unacylated ghrelin (UAG) affects glucose tolerance but has no effect on tumour volume or on other metabolic parameters. *Rag1^−/−^* mice fed a 23% high-fat diet (HFD) or low-fat diet (LFD) were injected with subcutaneous PC3 xenografts and administered UAG (100μg/kg/day, i.p.) (*n*=6 HFD, *n*=10 LFD) or PBS control (*n*=8 HFD, *n*=10 LFD) once tumours were palpable. Mean ± s.e.m. * *P*≤0.05. (A) Tumour volume (mm^3^) measured over time (*P*=0.57), (B) tumour volume (mm^3^) (*P*=0.55) and (C) tumour weight (g) measured at experimental endpoint were not significantly different between UAG- and PBS-treated mice fed HFD or LFD. Mean ± s.e.m. Mann-Whitney test. (D) Body weight (g) of mice at endpoint (*P*=0.08), (E) epididymal fat pad weight (g) (*P*=0.26) and (F) interscapular brown adipose tissue weight (g) (*P*=0.12) were not significantly different in UAG-treated mice compared to PBS-treated mice. Mean ± s.e.m. Mann-Whitney test. (G) HFD-fed UAG-treated mice (*n*=6) had significantly lower blood glucose 30 min post-glucose challenge compared to HFD-fed PBS treated mice (*n*=8), determined by intraperitoneal glucose tolerance test (IPGTT). Mean ± s.e.m. Two-way ANOVA. * *P*=0.025. (H) Fasting blood glucose (mM) (*P*=0.50) and (I) fasting blood insulin (ng/ml) were not altered in UAG-treated compared to PBS-treated mice on either diet. Mean ± s.e.m. *P*=0.90. Mann-Whitney test. (J) Insulin resistance (HOMA-IR) (*P*=0.70), (K) steady state β-cell function (HOMA%B) (*P*=0.22) and (L) insulin sensitivity (HOMA%S) (*P*=0.70) were not altered in UAG-treated compared to PBS-treated mice on either diet. Mean ± s.e.m. Mann-Whitney test.

## Discussion

It has recently been recognised that UAG can under some conditions act as a functional ghrelin inhibitor, reducing ghrelin-mediated increases in plasma glucose [22, 26, 28, 72] and lipid [27, 29]. As the ghrelin axis also plays a role in the progression of a number of endocrine-related cancers [5, 32–34], including prostate cancer [5, 43], we hypothesised that UAG may have beneficial effects in advanced prostate cancer associated with metabolic syndrome. To evaluate this hypothesis, we examined the effect of UAG on a prostate cancer cell line *in vivo*.

In our diet-induced hyperinsulinaemic *Rag1^−/−^* mouse model [69], we investigated the effect of supraphysiological systemic UAG treatment (100μg/kg/day) on metabolic parameters and PC3 prostate cancer xenograft growth. No differences in metabolic parameters (fasting blood glucose, fasting blood insulin, insulin resistance, steady-state β-cell function, and insulin sensitivity) were observed with UAG treatment in HFD-fed mice. Other studies have found that UAG prevents insulin resistance and hyperglycaemia in short-term HFD-fed mice [73], observations which may stem from the ability of UAG to cross the blood-brain barrier and oppose the central actions of ghrelin on energy homeostasis [74]. Furthermore, in human clinical trials UAG improved glucose and lipid metabolism in healthy [29] and diabetic patients [28]. In our study, a decrease in bodyweight, epididymal fat pad weight, and interscapular brown adipose tissue was observed in HFD-fed UAG treated mice but this difference was not statistically significant. UAG did significantly reduce blood glucose levels at 30 minutes post-glucose challenge in HFD, but not LFD-fed mice, however. This is similar to other studies, which have only found positive effects of UAG on glucose tolerance in obese patients [72]. Similarly, in clinical trials, AZP-531 (a cyclised, truncated analogue of UAG) improved food-related behaviour, waist circumference, and glucose tolerance in Prader-Willi syndrome patients, but had no effect on body weight [31]. AZP-531 also prevents HFD-induced weight gain, insulin resistance, and impairment of glucose tolerance in mice [27].

To the best of our knowledge, this is the first report on the effects of UAG on cancer cell line xenograft growth *in vivo*. While our study and others show somewhat promising effects of UAG treatment on metabolic parameters, systemic UAG administration had no effect on prostate tumour xenograft size in mice fed a low-fat or high-fat diet. While preliminary, our study suggests that UAG administration, or targeting of endocrine UAG, may have limited therapeutic potential for prostate cancer, in patients with and without symptoms of metabolic syndrome.

## Acknowledgments

This work was supported by the National Health and Medical Research Council Australia (grant no. 1002255 and 1059021 to PLJ, LKC, ACH, and IS), the Cancer Council Queensland (grant no. 1098565 to LKC, ACH, and IS), the Australian Research Council (grant no DP140100249 to LKC and ACH), a QUT Vice-Chancellor’s Senior Research Fellowship (to IS), the Movember Foundation and the Prostate Cancer Foundation of Australia through a Movember Revolutionary Team Award, the Australian Government Department of Health, and the Australian Prostate Cancer Research Center, Queensland (LKC, ACH, and CCN). The funders had no role in study design, data collection and analysis, decision to publish, or preparation of the manuscript.

